# Brain dynamics for confidence-weighted learning

**DOI:** 10.1101/769315

**Authors:** Florent Meyniel

## Abstract

Learning in a changing and uncertain environment is a difficult problem. A popular solution is to predict future observations and then use surprising outcomes to update those predictions. However, humans also have a sense of confidence that characterizes the precision of their predictions. Bayesian models use this confidence to regulate learning: for a given surprise, the update is smaller when confidence is higher. We explored the human brain dynamics sub-tending such a confidence-weighting using magneto-encephalography. During our volatile probability learning task, subjects’ confidence reports conformed with Bayesian inference. Several stimulus-evoked brain responses reflected surprise, and some of them were indeed further modulated by confidence. Confidence about predictions also modulated pupil-linked arousal and beta-range (15-30 Hz) oscillations, which in turn modulated specific stimulus-evoked surprise responses. Our results suggest thus that confidence about predictions modulates intrinsic properties of the brain state to amplify or dampen surprise responses evoked by discrepant observations.

## Introduction

Popular learning algorithms like predictive coding (Rao and Ballard, 1999; Shipp et al., 2013; Aitchison and Lengyel, 2017; Spratling, 2017) and the delta rule (Rescorla and Wagner, 1972; Sutton and Barto, 1998) posit that expectations (or equivalently here, predictions) play a key role in learning from a sequence of observations. Those algorithms, in their simplest form, consist in updating the quantity that is learned in proportion to the prediction error, which is the difference between the prediction and the actual observation. This solution is simple and yet efficient even in changing and uncertain environments (Sutton and Barto, 1998; Yu and Cohen, 2008; Heilbron and Meyniel, 2019). Other, more sophisticated algorithms exist (Behrens et al., 2007; Nassar et al., 2010; Wilson et al., 2013; Mathys et al., 2014; Ritz et al., 2018; Moens and Zénon, 2019), they may formalize the discrepancy between predictions and observations differently (prediction error, improbability, surprise) but they all have in common that this discrepancy is the driving force of learning.

Recently, another aspect of human learning has been put forward: learning is accompanied by a sense of confidence about predictions. Interestingly, this sense of confidence follows, at least in part, the optimal principles of probabilistic inference; in that sense, it is rational (Nassar et al., 2010; Meyniel et al., 2015a; Boldt et al., 2019). Here, we embrace the proposal that this sense of confidence in a learning context indeed plays a functional role in learning (Nassar et al., 2010; Iglesias et al., 2013; Meyniel et al., 2015b; Meyniel and Dehaene, 2017), as prescribed by the optimal rules of probabilistic inference. Bayesian models, which obey those rules, show this property. In such models, the update is not only guided by discrepant observations, it is also regulated by confidence about predictions: for a given discrepancy, the update is smaller when the confidence associated with the prediction was larger. This confidence-weighting principle is not specific to learning, it is generally applicable whenever several sources of information must be combined (Ernst and Banks, 2002; Ma et al., 2006; Bang et al., 2014, 2017; Meyniel et al., 2015b; Rohe et al., 2019). In a learning context, confidence should set the balance between predictions and new data.

The notion of confidence-weighting is related to others, like selective attention (Dayan et al., 2000; Meyniel et al., 2015b), the weight of evidence (Rohe et al., 2019), the precision of predictions (Iglesias et al., 2013; Mathys et al., 2014; Vossel et al., 2014) discussed at the end of this article. Here, we propose to use optimal Bayesian models as a benchmark to formalize, at a computational level, the learning process. In particular, we formalize the notion of discrepancy between predictions and observations with the information-theoretic measure of surprise (Shannon, 1948; Strange et al., 2005; O’Reilly et al., 2013), and the notion of confidence about a prediction, as the precision of the posterior predictive distribution (Summerfield et al., 2011; Yeung and Summerfield, 2012; Meyniel et al., 2015a; Boldt et al., 2019).

Previous studies reported conflicting results about the existence of a confidence-weighting mechanism in the brain (Jepma et al., 2016; Meyniel and Dehaene, 2017; Nassar et al., 2019). Here, we used a probability learning task previously developed, in which the participants’ reports of probability estimates and the associated confidence levels are compatible with optimal Bayesian inference. Our goal is two-fold. Numerous studies reported the existence of surprise signals in the brain, i.e. neural responses that are more vigorous for unexpected stimuli (Hillyard et al., 1971; Squires et al., 1976; Nassar et al., 2012; Wacongne et al., 2012; Kolossa et al., 2013; O’Reilly et al., 2013; Summerfield and de Lange, 2014; Garrido et al., 2016; Heilbron and Chait, 2018; Maheu et al., 2019). We will test whether surprise responses show an additional effect of confidence as prescribed by the confidence-weighting principle, making them suitable for close-to-optimal updates. Second, previous studies suggested that evoked responses are gated by the state of brain networks, which is characterized by specific synchronization of oscillatory activity (Hipp et al., 2011; Siegel et al., 2012; Baumgarten et al., 2016; Hahn et al., 2019; Iemi et al., 2019) and controlled by neuromodulators (Aston-Jones and Cohen, 2005; Reimer et al., 2014; Safaai et al., 2015; Vazey et al., 2018; Zaldivar et al., 2018; Rodenkirch et al., 2019). We will test whether confidence modulates the spectral properties of brain activity and the neuromodulatory state reflected by non-luminance related changes in pupil size.

## Results

### An optimal framework to study confidence in a learning context

We adopted a probability learning task (**Fig. 1A**) that is amenable to Bayesian modeling, and in which the human inference is well accounted for by Bayesian models. Subjects were presented with auditory sequences made of two tones (say A and B), presented randomly according to predefined transition probabilities between successive tones: p(A|A) and p(B|B). Note that p(B|A) and p(A|B) are deduced from the two other quantities as 1-p(A|A) and 1-p(B|B). Those transition probabilities changed abruptly and unpredictably in the course of the experiment, at “change points”. The generative model of the task therefore has three levels, organized hierarchically: 1) the tones, 2) the transition probabilities governing the tones and 3) the probability of change point, which was fixed across trials. We used Bayesian inference to invert the generative process of the task **(Fig 1B)** and estimate, at any given trial, the transition probabilities currently generating the tones, given previous observations and knowledge of the actual task structure. This inference returns full posterior distributions for the transition probabilities p(A|A) and p(B|B), see **Fig 1C**.

**Figure 1:**
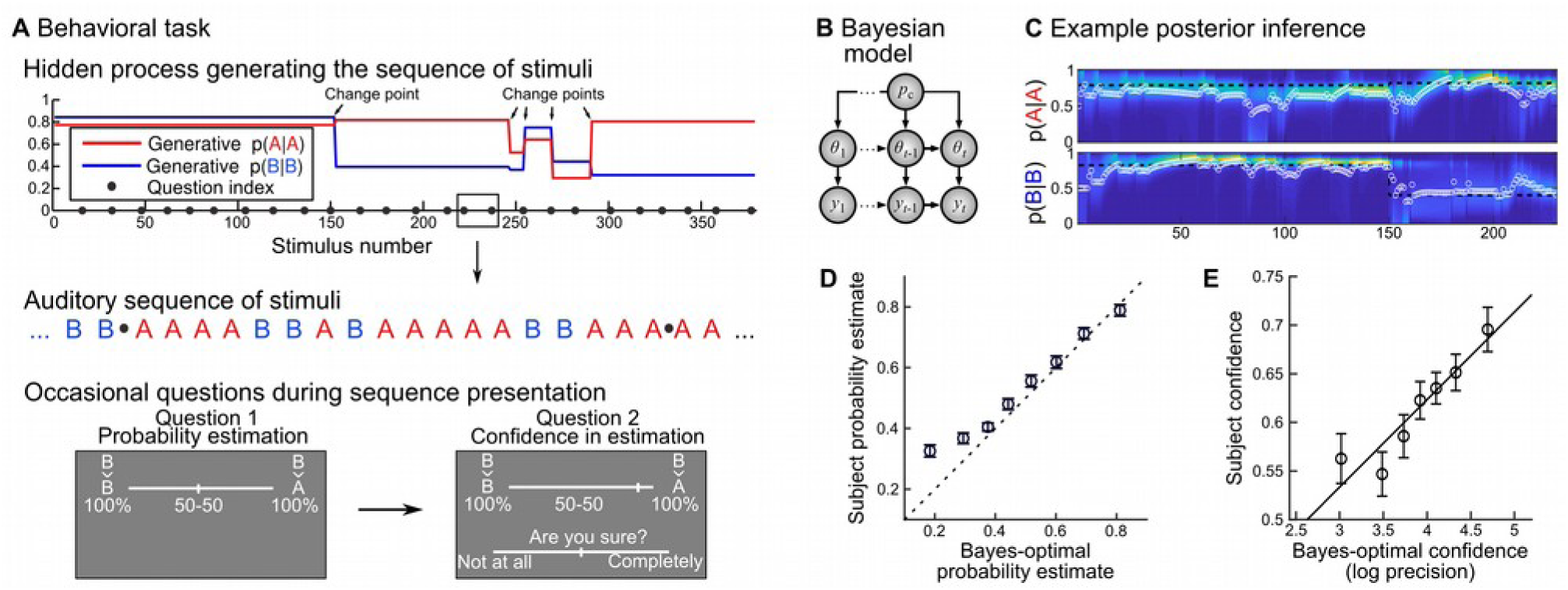
Rational confidence levels during a probability learning task. **(A)** Subjects listened to sequences of two tones (denoted A and B) generated randomly following (order 1) transition probabilities that changed abruptly at unpredictable moments. Subjects knew about this generative process and the tones were perceived without ambiguity. Occasionally subjects were asked to report the probability of the next tone, which amounts to reporting the transition probability relevant at the moment of the question, and then their confidence about this probability estimate. They used continuous sliders for both reports. (**B)** We design a Bayes-optimal, hierarchical model that corresponds to the generative process used in the experiment: given current and past observations *y*s, it estimates the momentary transition probabilities *θ*_*t*_, knowing that there is a fixed probability p_c_ (=1/75) of a change point at every trial. (**C)** The Bayes-optimal, hierarchical model returns a posterior, time-varying, two-dimensional distribution, each dimension corresponding to a transition probability. This distribution evolves as a function of the observations received; the heat-map shows the evolving distribution, color-coded and stacked horizontally (a vertical slice corresponds to the distribution at a given moment). We formalized the answers to question 1 (probability report) and 2 (confidence) as the mean of the distribution (white circle) and its log-precision, respectively. (**D-E)** Subjects reports are plotted against bins of Bayes-optimal values, as mean ± s.e.m.

Subjects, as the Bayes-optimal model, were fully informed about the task structure; the only difference is that they were not given the numeric value of the frequency of change points, but the qualitative indication that they are rare. We probed their inference occasionally by asking them to report the probability of the next tone (question #1), and then their confidence about this estimate (question #2). In the Bayes-optimal model, the answer to question #1 corresponds to the mean of the posterior distribution of the relevant transition probability (the one that corresponds to the tone presented on the previous trial), and we formalized the answer to question #2 (confidence) as the log-precision of this distribution (see Methods) because when precision is high, the posterior evidence is highly concentrated around the mean, which gives credence to this value (**Fig 1C**).

Subjects’ probability estimates were highly correlated with the Bayes-optimal solution (**Fig 1D**): Pearson ρ=0.62 ± 0.04 sem, t_23_=16.75, p=2.2 10^-14^. The same was true for confidence report, although to a lesser extend (**Fig 1E**): ρ=0.23 ± 0.04 s.e.m., t_23_=6.19, p=2.6 10^-6^. In this study, it is import to distinguish prediction from confidence, since we test their different roles in the learning process. We thus tested that subjective confidence is not reducible to the prediction itself or related quantities. We estimated a multiple regression model of subjective confidence, using as predictors: the Bayes-optimal prediction about the next tone, the corresponding predictability (formally, the expected surprise or equivalently, the entropy of prediction), the surprise from the previous trial, and the prediction error on the previous trial, as well as the subject’s prediction about the next tone and the corresponding predictability (we cannot include subjective surprise and prediction error from the previous trial since we do not know them due to the occasional nature of the questions). An effect of some of those factors on confidence exists in the normative model itself (Meyniel et al., 2015a) and was significant in subjects (effect of Bayes-optimal previous surprise: p=0.031; entropy of subject’s prediction: p=3.1 10^-8^), but importantly, the residuals of the multiple regression remain correlated with Bayes-optimal confidence (β=0.039 ± 0.011, t_23_=3.5, p=1.9 10^-3^,) indicating that subjective confidence cannot be reduced entirely to those factors.

Below, we leverage several features of this task. First, the Bayes-optimal model provides an account of the subjects’ probability estimates and confidence on the question trials, which licenses the use of this model to study the brain mechanisms of inference during the no question trials. The no question trials are advantageously numerous and unperturbed by motor artifacts or other processes related to answering questions. Second, the use of change points induce trial-to-trial variations in confidence, which facilitates the exploration of the neural correlates of confidence. The use of transition probabilities further increases those variations by de-correlating confidence from one trial to the next (see **Fig S1** for an example).

### Evidence for a confidence-weighting of surprise in evoked responses

Many brain responses evoked by stimuli are known to be modulated by surprise: they are more vigorous for unexpected stimuli. Here, we tested whether we could identify such surprise responses and whether they would show additional effects of confidence following the confidence-weighting principle, i.e. dampened responses for higher confidence. To this end, we estimated a multiple regression model, systematically and independently for all sensors and peri-stimulus times, from −0.1 to 0.8 s. Note that the inter-stimulus interval is 1.4 s, such that this time window only contains the current stimulus. We included three factors: the stimulus identity (coded as a binary variable), the Bayes-optimal surprise to the current stimulus and the Bayes-optimal confidence about the prediction of the current stimulus identity. This analysis yielded significant results for the three factors (cluster-forming p<0.001, cluster-level p<0.05, two-tailed, n=1,000).

We also performed a more conservative analysis, yielding very similar results. Confidence being correlated with predictability in the Bayes-optimal model (ρ=-0.37 ± 0.023 s.e.m., t_23_=-16.3, p=4.2 10^-14^), the effect of confidence we found could be due to correlation with the prediction itself. We estimated another regression model, replacing the Bayes-optimal confidence with the residual confidence, after linearly regressing out effects of prediction and predictability (see Methods). **Fig 2A** shows the result of this analysis and various significant effects (cluster-forming p<0.001, cluster-level p<0.05, two-tailed, n=1,000): the stimulus identity (105-160 ms) co-occuring with a first surprise response (80-155 ms) followed by a second one (160-290 ms) and a late, prolonged one (360-715 ms). Importantly, an effect of confidence (155-230 ms) overlapped in space and time with the effect of surprise. The time-course activity in the confidence cluster (**Fig 2B**) shows that the response around 200 ms, characterized by an inward field (negative sign), is more vigorous for unexpected stimuli, and dampened for higher confidence, akin to the confidence-weighted surprise signal predicted by Bayesian learning. Source reconstruction within 155-230 ms (**Fig 2C**) revealed that the negative effect of confidence and the positive effect of surprise on brain activity overlapped notably in the right inferior frontal sulcus and intermediate precentral sulcus, rather than in the primary auditory cortices.

**Figure 2:**
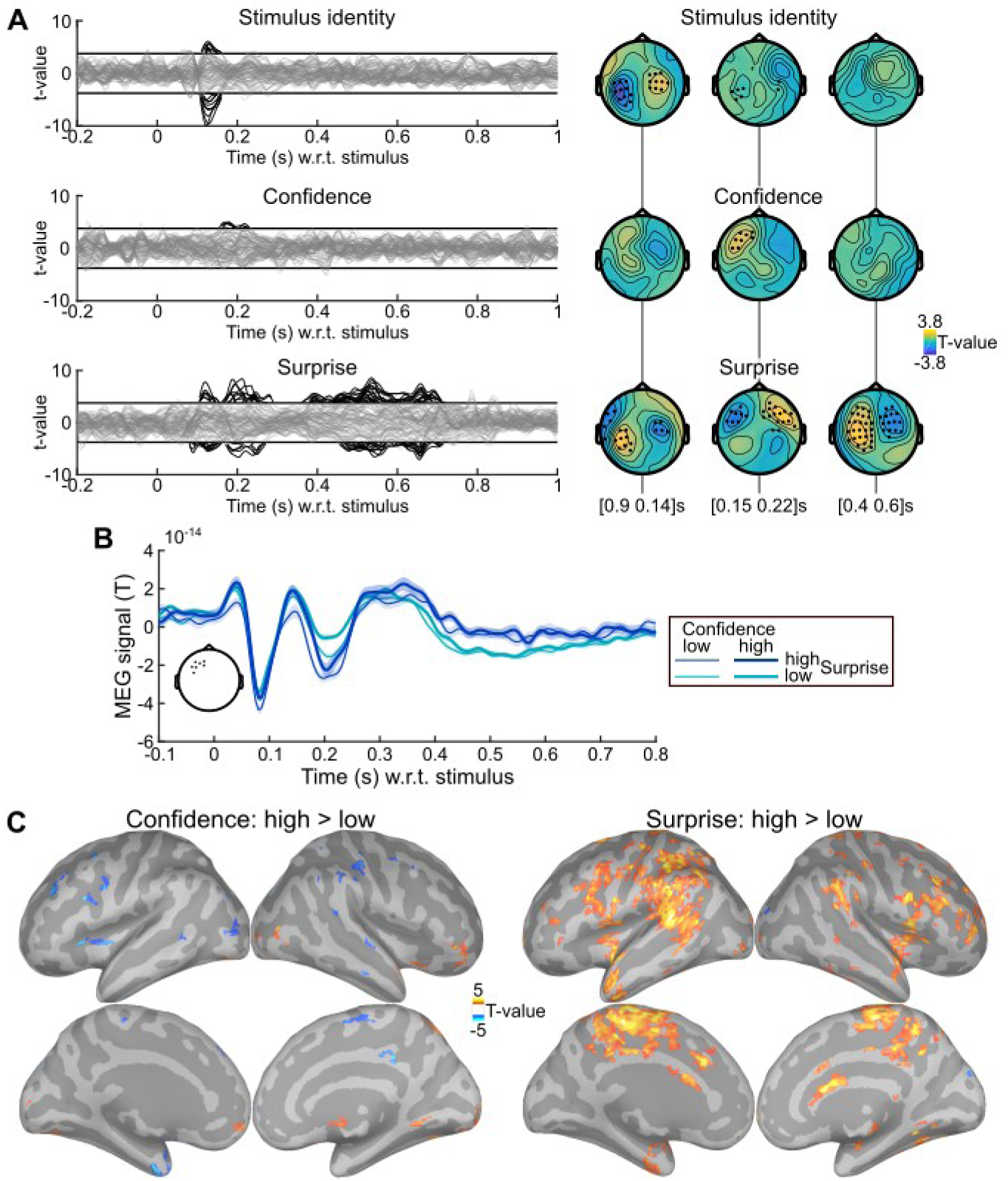
Evidence for surprise and confidence-weighted surprise in evoked activity. **(A)** The stimulus-locked evoked response was analyzed with mass-univariate regression. The value at each sensor and peri-stimulus time was regressed, across trials, using a multiple linear model with three factors: the stimulus identity, the (residual) Bayes-optimal confidence about the probability of the current stimulus and the Bayes-optimal surprise (i.e. negative log likelihood). The time series show the significance, as t-values, of regression coefficients, with one line per sensor; topographies show the average t-value in three time windows. Highlighted time points and sensors survive correction for multiple comparisons across peri-stimulus times and sensors (cluster-forming p<0.001 two-tailed, cluster-level p<0.05 two-tailed, n=1,000). (**B)** Time series of MEG signal (mean ± s.e.m. estimated after subtracting the subject-mean) in the cluster of sensors (see inset) showing a significant effect of Bayes-optimal confidence. Trials were sorted into high and low surprise (with respect to 1 bit of surprise, i.e. observation likelihood of 0.5) and high and low confidence (median split). The response around 200 ms is more extreme for higher surprise and less extreme for higher confidence. (**C)** Source reconstruction was computed within 155-230 ms for trials with high and low confidence (median split), high and low surprise (median split). The significance of the paired difference between high and low levels is shown separately for confidence and surprise (threshold: p<0.05, two-tailed, minimum cluster size: 50 vertices, total vertices: 306,716). Increased signal for higher surprise and lower signal for higher confidence is found notably in the right inferior frontal sulcus and intermediate precentral sulcus.

We further tested the robustness of the confidence effect found within 155-230 ms in the significant sensors shown in **Fig 2B**. First, the regression model did not include an interaction between surprise and confidence, because their effects on model update is theoretically mostly additive (keep in mind that those quantities are on a log-scale). Their interaction, when included in the previous linear regression model, was not significant (t_23_=0.29, p=0.77) and left the results unchanged. Second, the temporal profiles of Bayes-optimal confidence share commonalities across subjects, for instance, it is low at the beginning of a session, and then waxes and wanes multiple times (see **Fig S1**). One concern is that those temporal characteristics may drive, for spurious reasons, the correlation between confidence and the MEG signal. To rule out this possibility, we shuffled the time-courses of (residual Bayes-optimal) confidence with respect to MEG data across subjects to estimate a null distribution for the correlation between confidence and MEG signal, and a Z-statistics. This approach controls for the temporal characteristics of Bayes-optimal confidence observed across all sequences presented to subjects, it is thus very conservative; yet the correlation between confidence and MEG signal remained significant (z=0.95 ± 0.19 s.e.m., t_23_=4.92, p=5.7 10^-5^).

### Modulation of brain states by confidence: spectral components

Next, we examined whether Bayes-optimal confidence would correlate with spectral components of the MEG signal, since power in low frequencies (<40 Hz) typically characterizes the state of large networks, which would then respond differently to incoming stimuli, thereby implementing a confidence-weighting mechanism for surprise (see Discussion). The signal was decomposed across peri-stimulus times and frequencies (6 – 40 Hz) and analyzed from −0.5 to 0.9 s. We used the same regression model as used for evoked response, with stimulus identity, Bayes-optimal surprise and residual Bayes-optimal confidence. We found significant effects (cluster-forming p<0.001 and cluster-level p<0.01 two-tailed, corrected for multiple comparisons, n=2,000) of surprise and confidence which were all positive: the higher the confidence (or surprise), the higher the power. The surprise effect, given its topography, timing and low frequency simply reflected the evoked responses identified in **Fig 1A**. In contrast, the confidence effect was much more extended in terms of time, frequency and sensor. For simplicity, we grouped significant clusters in pre and post-stimulus clusters, and further sub-divided the latter in alpha and beta range, with respect to 12 Hz (**Fig 2A**). Note that the confidence effect was present even before the stimulus, see for illustration the time courses (**Fig 2B**) and topography (**Fig 2C**) of the confidence effect in the 15-25 Hz band.

We further tested the robustness of those confidence effects in those three clusters. All clusters passed the tests, but showed that the effect was most robust in the post-stimulus, beta-band cluster; therefore, from now on we focus on this one (unless otherwise specified). First, we controlled that the correlation we found was not driven by the general temporal profile of Bayes-optimal confidence in the course of experimental sessions, using the same permutation procedure as described for the evoked responses. The effect remained significant (z=1.16 ± 0.16 s.e.m., t_23_=7.35, p=1.8 10^-7^). We also tested whether the effect survived the inclusion of confidence from the previous trial in the regression model, which was true: β=2.5 10^-26^ ± 4.3 10^-27^ s.e.m., t_23_=5.93, p=4.8 10^-6^; this test is particularly relevant for the pre-stimulus cluster (β=6.8 10^-26^ ± 1.8 10^-26^ s.e.m., t_23_=3.85, p=8.2 10^-4^), denoting a preparation of brain networks to the upcoming stimulus based on the confidence hold specifically in the prediction. This effect suggests that power can change from one trial to the next, following changes in confidence. We tested this possibility by considering pairs of consecutive trials, sorting them into low and high confidence (median split) on the current trial and further sorting them into high vs. low confidence (median split) on the next trial. High-to-low and low-to-high transitions in Bayes-optimal confidence indeed corresponded respectively to decreases and increases in power from one trial to the next, see **Fig S2**.

If some spectral properties of neural signals correlate with the (Bayes-optimal) confidence about the prediction even before the presentation of the next stimulus, then they may be predictive of the confidence reported by subjects when the stimulus is actually replaced by a question. We trained a ridge regression model to predict the subject’s confidence report based on the pre-stimulus power (average between −500 and 0 s with respect to the omitted stimulus) in frequencies ranging from 6 to 40 Hz, and evaluated its predictive accuracy in a cross-validated manner (see Methods). Out-of-sample predictions indeed co-varied weakly but significantly with subjective confidence (Pearson ρ=0.06 ± 0.02 s.e.m., t_23_=2.43, p=0.023, **Fig 3D**). To explore the frequencies contributing to this predictive power, we estimated the ridge regression weights on the full dataset of each subject (see **Fig 3F**). Those weights must be interpreted with caution because they depend on the covariance of the data, and may be different across subjects; nevertheless they suggest a positive effect of (low) beta range power (15-20 Hz) similar to the effect of Bayes-optimal confidence during the no-question trials. However, power in the alpha range (around 10 Hz) was negatively related to subjective confidence, unlike the effect of optimal confidence on power during no-question trials, suggesting the existence of multiple processings behind subjective confidence.

**Figure 3:**
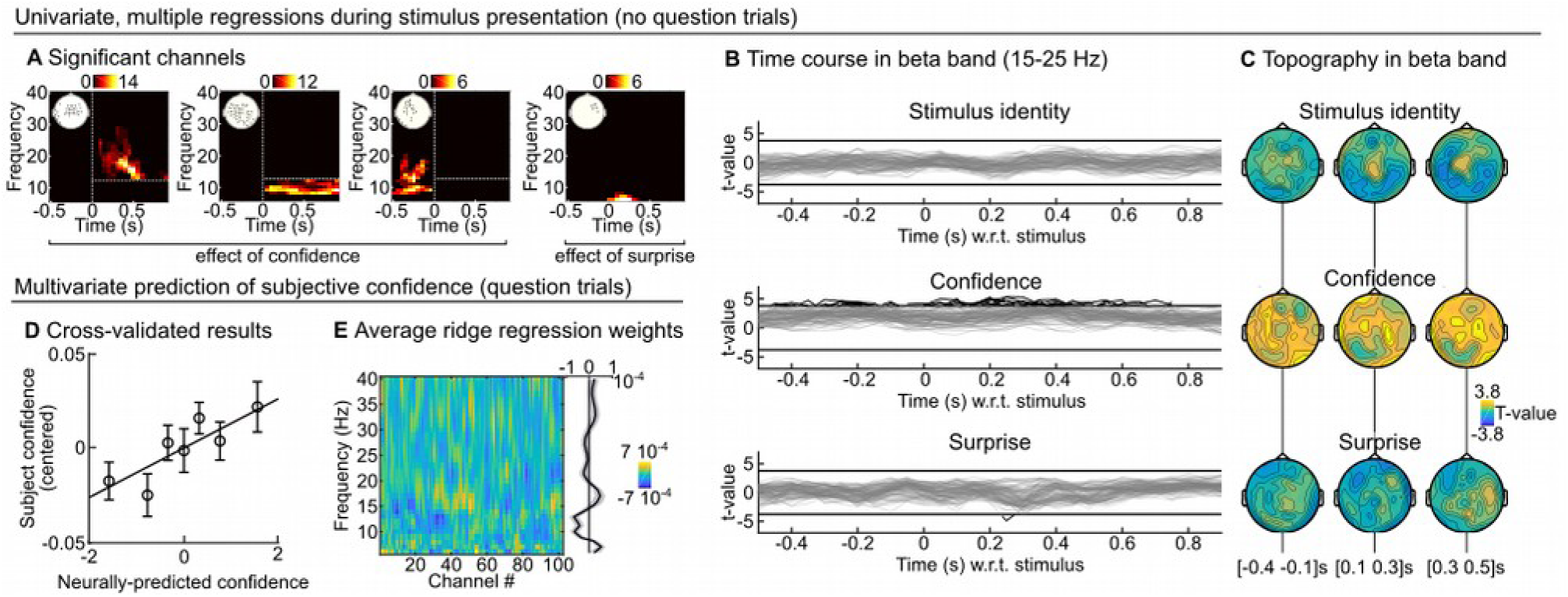
Beta-range oscillations reflect optimal and subjective confidence. **(A)** The peri-stimulus power was analyzed with mass-univariate regression. The value at each sensor, peri-stimulus time and frequency was regressed, across trials, using a multiple linear model with three factors: the stimulus identity, the (residual) Bayes-optimal confidence about the probability of the current stimulus and the Bayes-optimal surprise. We used correction for multiple comparisons across peri-stimulus times, sensors and frequencies (cluster-forming p<0.001, cluster-level p<0.01 two-tailed, n=2,000). Several clusters were found for Bayes-optimal confidence, which were regrouped for clarity as: post-stimulus effect in beta band (>12 Hz); post-stimulus effect in the alpha band (≤12 Hz) and pre-stimulus effect in alpha-beta band. For each cluster, the inset shows the significant sensors and the heat-map shows the number of significant sensors for each time-frequency pair. **(B-C)** For illustration, we show the significance levels (as t-value) of each predictor variable in the 15-25 Hz band, as time courses (**B**, one line per sensor; the threshold correspond to p<0.001 two-tailed, n=2,000) and topographies (**C**). **(D-E)** During question trials, the stimulus was replaced by a question, asking the subject for her prediction about the stimulus identify. We trained a multivariate (ridge) regression model in a cross-validated manner to predict, based on the pre-stimulus power (average within −500 to 0 ms relative to question) across frequencies, the confidence report of the subject. **D** shows the cross-validated prediction accuracy (mean ± s.e.m.) and **E** shows the (non cross-validated) regression weights for each frequency and channels (together with the average across channels).

### Modulation of brain states by confidence: pupil-linked arousal

The state of brain networks is also modulated by neuromodulation, in particular the arousal state that is reflected in changes in pupil diameter. Subjects were fixating and the task was auditory, without visual stimuli or change in luminance, the pupil diameter therefore reflected here the subject’s internal state. We distinguished two aspects of the pupil diameter: phasic and tonic levels, which we measured respectively as changes relative to a pre-stimulus baseline (−250 to 0 ms) and absolute changes. The phasic responses showed a transient increase in response to the stimulus (**Fig 4A**); we used a multiple linear regression for each peri-stimulus time points, including the Bayes-optimal surprise and Bayes-optimal confidence. The phasic levels showed only an effect of Bayes-optimal surprise whereas the tonic levels showed only an effect of Bayes optimal confidence (cluster-forming p<0.05; cluster-level p<0.001, two-tailed, n=10,000). In order to test the robustness of this effects, we repeated the same multiple regression analysis but considering now the residual Bayes-optimal confidence (cluster-level p=0.005, n=10,000) and to control for the temporal profile of Bayes-optimal confidence, we used the same permutation analysis as described above (cluster-level p=0.03 one-tailed, n=10,000).

**Figure 4:**
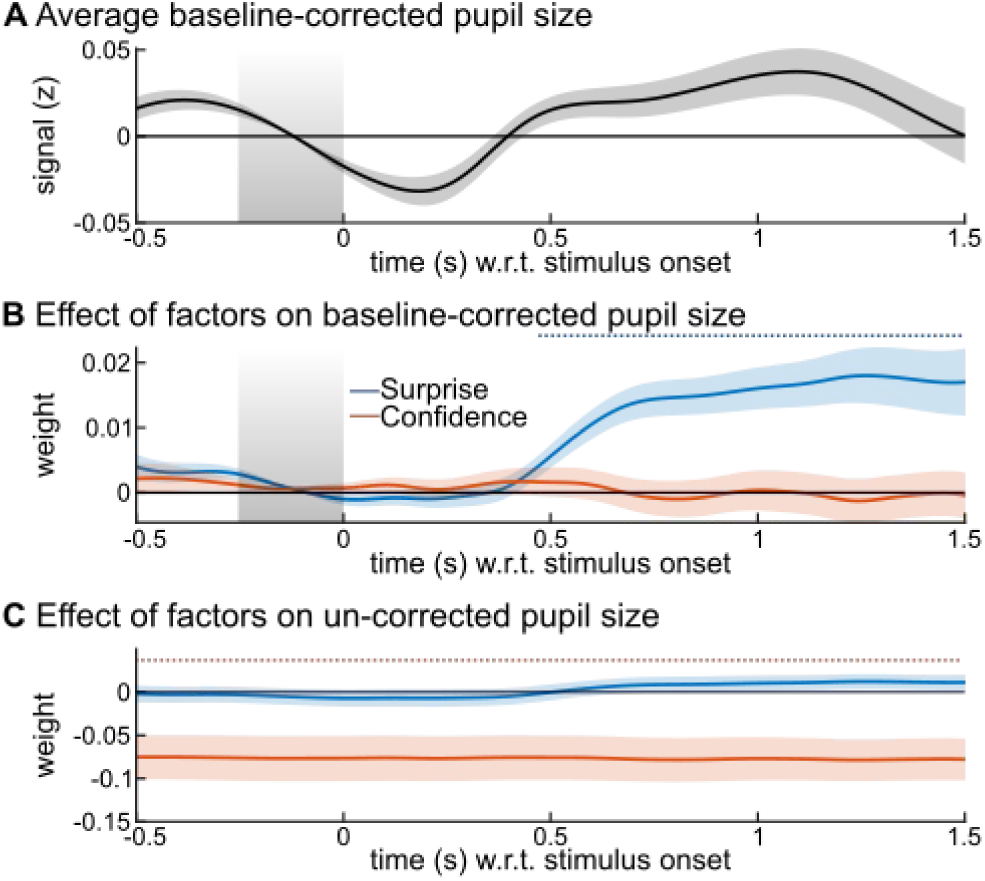
Tonic and phasic pupil size distinctly reflect confidence and surprise. The z-scored pupil signal was analyzed with mass-univariate regression. The value at each peri-stimulus time was regressed, across trials, using a multiple linear model with three factors: the (residual) Bayes-optimal confidence about the probability of the current stimulus, the Bayes-optimal surprise. We used the baseline (see grey area) corrected pupil size as a measure of phasic response. **(A)** shows its average values and **(B)** the effects of factors. We used the uncorrected pupil size as a measure of its tonic level; **(C)** shows the effect of factors. Time-series show mean ± s.e.m.; the horizontal dashed lines indicate significant time points, corrected for multiple comparisons (cluster-forming p<0.05; cluster-level p<0.05, two-tailed, n=10,000).

### Modulation of evoked responses by brain states

The above analyses showed that the confidence about predictions modulates brain states, in particular large-scale beta-range oscillations and the arousal state indexed by tonic pupil size. If those brain states play a role in the confidence-weighting of surprise, their level should modulate the confidence-weighted surprise responses evidenced in **Fig 2**, around 200 ms. In particular, they may capture trial-by-trial variations in the subject-specific states and thus account for the dampening of this surprise response on top of the effect predicted by the Bayes-optimal confidence. In order to test this possibility, we estimated two multiple regression models onto the evoked responses shown in **Fig 2B**, including the Bayes-optimal surprise, Bayes-optimal (residual) confidence, and the trial-by-trial value of either the post-stimulus beta-band activity (using the cluster shown in **Fig 2A**) or the tonic pupil size (averaged within −250 to 0 ms relative to stimulus onset). Bins of successive trials were used to achieve more robust results, see Methods. We corrected results for multiple comparisons using a cluster-forming p<0.05 and cluster-level p<0.05, one-tailed, n=10,000 (**Fig 5**).

**Figure 5:**
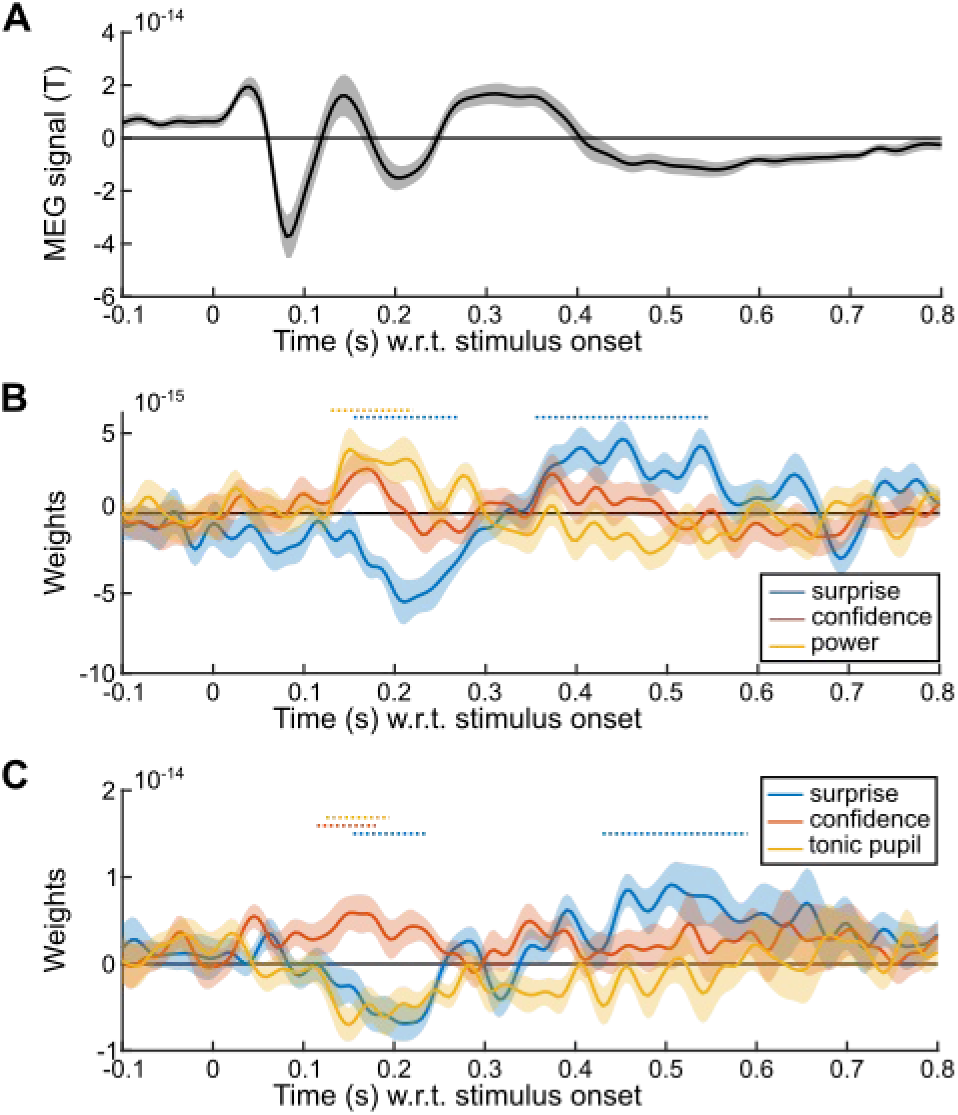
Tonic pupil size and beta-range oscillations shape the evoked response. The signal in the confidence cluster shown in Fig 2B was analyzed with mass-univariate regression. **(A)** Average MEG signal in the confidence cluster shown in Fig 2B. The sign of the signal is useful to interpret the effect of factors. **(B)** Results of a multiple regression model comprising: the (residual) Bayes-optimal confidence about the probability of the current stimulus, the Bayes-optimal surprise and the power in the post-stimulus, beta band cluster (see Fig. 3A). **(C)** Results of the same multiple regression model but with tonic pupil size instead of beta-band power. Time-series show mean ± s.e.m., horizontal dashed lines indicate significance corrected for multiple comparisons (cluster-forming p<0.05; cluster-level p<0.05, one-tailed, n=10,000).

This analysis replicates the result found in Fig 2: an effect of Bayes-optimal surprise around 200 ms (as well as later) together with an opposite effect of Bayes-optimal confidence. More beta-band power being correlated with higher confidence, beta-band power should dampen the surprise response, whose sign was negative (corresponding to an inward field), this negative effect (dampening) onto a negative surprise response (inward field) should result in a positive regression coefficient, a result that we observed and which was significant only around 200 ms. By contrast, higher tonic pupil levels correlated with lower confidence, tonic pupil should therefore strengthen the surprise response, which was negative (inward field), an effect that should result in a negative regression coefficient, which we observed significantly and selectively around 200 ms.

In order to better interpret those effects, it is important to know whether those beta-band oscillations and the tonic pupil size are two sides of the same coin (but note that pupil data is available only in 18/24 subjects, reducing statistical power). If so, one would expect a negative trial-to-trial correlation, since those signals co-vary, respectively, positively and negatively with Bayes-optimal confidence. However, this correlation was negligible, if not even slightly positive (Pearson ρ=0.043 ± 0.03 s.e.m., t_17_=1.53, p=0.14). Instead of using separate multiple regression models, one can include those two signals in the same model of the evoked response, along with the Bayes-optimal surprise and confidence. The regression coefficients become difficult to interpret since the predictors (pupil size, beta-band power) share correlation with confidence, which itself correlates with the evoked response. However, we found a significant effect of pupil from 115 to 220 ms (cluster-forming p<0.05 and cluster-level p<0.05, one-tailed, n=10,000).

## Discussion(1500)

Subjects estimated the probabilities that characterize a sequence of inputs, and the confidence associated with those estimates, following (to some extent) the principles of Bayesian inference. Those estimates were accurate despite the presence of volatility. Bayesian inference leverages the confidence-weighting principle to remain accurate in the face of volatility, by dynamically modulating the update prompted by surprising observations. The MEG responses evoked by stimuli showed numerous signatures of surprise (vigorous responses to unexpected stimuli) and for one of them, an additional effect of confidence, conforming to the confidence-weighting principle. Whole-brain beta-band power increased with optimal confidence and predictive of the actual subject’s confidence. Lack of confidence increased the tonic pupil size while phasic dilation reflected surprise. Desynchronization of beta-band power and higher tonic pupil size increased the evoked, confidence-weighted surprise response. Overall, the results indicate that changes in brain states indexed by beta-range power and neuromodulation, are related to differential responses to surprising stimuli, providing a mechanism for a dynamic confidence-weighting of learning.

We now discuss those results in the context of previous studies, which requires to delineate precisely what aspects of confidence we investigated here. It is the confidence that accompanies probability estimates on a trial-by-trial basis, thus it is not “global” or related self-confidence (Rouault et al., 2019). It is also not related to whether the stimulus is perceived clearly or ambiguously, which is important in general for inference (Kepecs et al., 2008; Mathys et al., 2014; Meyniel et al., 2015b; Pouget et al., 2016; Bang and Fleming, 2018) but does not play a role here since stimuli were perceived without ambiguity. In addition, most results here are based on Bayes-optimal confidence: the one that the subject *should* hold. It was indeed correlated with subject’s confidence (the confidence actually hold by subjects) and beta-band power (which itself was also predictive of subject’s confidence). Yet, Bayes-optimal confidence remains only an imperfect model of subject’s confidence and both quantities, although related here, should not be conflated.

The type of confidence studied here (about a learned estimate) is also different from most studies, which focused on confidence about memory (Koriat, 2012; Fleming et al., 2014) and confidence about decisions (Kepecs and Mainen, 2012; Meyniel et al., 2015b; Pouget et al., 2016). Those different types of confidence correspond theoretically to different constructs: confidence about a memory or decisions correspond to the probability of the decision or memory being correct (Pouget et al., 2016; Fleming and Daw, 2017), whereas confidence about a learned estimate (even when this estimate is itself a probability) corresponds to a higher-order quantity: the precision of a posterior distribution (Yeung and Summerfield, 2012; Meyniel et al., 2015a; Navajas et al., 2017; Boldt et al., 2019).

Last, previous studies have not always disentangled confidence from related quantities. In many situations, confidence about predictions (i.e. posterior precision) is related to predictability. Previous studies showed that the precision of predictions modulates reaction times (Vossel et al., 2014), functional magnetic resonance imaging (fMRI) activity in fronto-parietal networks (Vossel et al., 2015), pupil size (Vincent et al., 2019), but with tasks and analyses that leave unclear whether the effect is genuinely about precision, or also about predictability. A similar and related confound exists between expectation and attention, which are often related in practice but distinct in theory (Summerfield and de Lange, 2014). Here, we controlled specifically for this correlation by using the *residual* confidence in our analyses. This is important because many brain signals are associated with predictability (Friedman et al., 2001; Strange et al., 2005; Summerfield and de Lange, 2014; Vossel et al., 2015; Kok et al., 2017; Vincent et al., 2019).

We now discuss more broadly the putative mechanisms of confidence-weighting in a learning context. Our interpretation is that confidence modulates the state of brain networks, which then process the feedforward input differently, in particular by increasing (low confidence) or decreasing (high confidence) the response to surprising stimuli, thereby modulating the effect of surprise onto updating. This interpretation is in line with the notion of neural gain: when the neural gain is higher (low confidence), expected and unexpected stimuli, for the same difference in expectation, elicit a larger difference in response. The neural gain is largely thought to depend on attention, neuromodulation and oscillatory activity in brain networks, three factors that we discuss below.

The role of confidence on update described here is computationally identical to the one ascribed to selective attention: some observations are given more weight than others; this link has been already discussed by others (Dayan et al., 2000). This is also related to the notion of neural gain which enhances or suppresses the effect of a given stimulus on further, downstream processing (Eldar et al., 2013). Noradrenaline was proposed as a major modulator of the neural gain (Aston-Jones and Cohen, 2005). This is particularly interesting since we found modulations of tonic pupil size by confidence. Specifically, we found larger pupil size for lower confidence, which is consistent with previous studies (Colizoli et al., 2018). Non-luminance based changes in pupil size reflect the arousal state and in particular, noradrenaline release. Correlation between pupil size and firing activity in the locus coeruleus, the main nucleus releasing noradreline in the brain (Salgado et al., 2016), was found in both rodents (Reimer et al., 2014) and macaque monkeys (Joshi et al., 2016). In addition, noradrenergic activity, unlike cholinergic activity, correlates with both fast and slow changes in pupil size (Reimer et al., 2016). The effect of confidence on tonic pupil found here could therefore arise from a change in noradrenergic activity. This noradrenergic activity changes neurons’ membrane potential (McGinley et al., 2015) and its slow fluctuations (Reimer et al., 2014), promoting the selectivity of sensory processing, akin to the neural gain model. Similarly, activation of the locus-coeruleus (and increased pupil size) promotes feature selectivity in the sensory domain (Krishnamurthy et al., 2017; Rodenkirch et al., 2019). Changes in pupil size are also related to changes in the processing of sensory information and performance during perceptual decision making tasks (de Gee et al., 2014, 2017), with different effects of phasic and tonic pupil size (van Kempen et al., 2019) as in the present results.

In line with this noradrenergic modulation of neural gain, previous studies showed that larger tonic pupil size renders fMRI responses more extreme, i.e. lower or higher (Eldar et al., 2013). Larger pupil size and higher activity of the locus coeruleus are also associated with better memorization (Hoffing and Seitz, 2015; Takeuchi et al., 2016), and thus a long lasting effect of the current observations. Rodent studies showed that one-shot learning can occur with noradrenergic input to the hippocampus (Wagatsuma et al., 2018). More generally, a role for noradreline in confidence-weighted learning is consistent with the fact that increased pupil size co-occurs with increased updating (Nassar et al., 2012) and reset of current strategies (Devauges and Sara, 1990), and that more volatile learning contexts are associated with larger pupil size (Pulcu and Browning, 2017; Vincent et al., 2019). Pharmacological studies in humans also provide supportive evidence, showing that manipulation of noradrenergic activity impacts the estimation of confidence in a decision task (Hauser et al., 2017), learning dynamics when unexpected changes occur (Marshall et al., 2016) and the effect of surprise on update (Jepma et al., 2016, 2018).

The modulation of responses to new observations (notably surprising ones) by the current state of neural network has also been associated repeatedly to specific rhythms of network activity, which provide signatures of networks’ dynamics and computations (Wang, 2010; Siegel et al., 2012). This state-dependent modulation should aim at prioritizing new evidence when confidence is low, and on the contrary, preserve current estimates from conflicting evidence when confidence is high. Our data indicate that beta (and perhaps alpha) band power could play a key role in such a prioritization. This result is in line with the previous proposal that higher beta-band power preserves the current state of networks, promoting a “status quo” rather than a change (Engel and Fries, 2010) which may also be true for lower (alpha) frequencies (Klimesch et al., 2007). Beta-band power in particular is associated with feedback signals (Bastos et al., 2015), stronger attention and top-down control (Buschman and Miller, 2007; Salazar et al., 2012), and the prioritization of the current brain states over new inputs (Spitzer and Haegens, 2017). In support to this view, stronger low frequency (<30 Hz) oscillations attenuate early evoked sensory responses (Iemi et al., 2019), increase the criterion of perceptual detection (resulting in fewer detections) by modulating baseline excitability (Limbach and Corballis, 2016; Benwell et al., 2017; Craddock et al., 2017; Iemi et al., 2017) and degrade performance in perceptual decisions (Haegens et al., 2011).

It is noticeable that the modulation of evoked surprise responses was confined to early post-stimulus latencies (around 200 ms) rather than occurring later, as could be expected for instance from the proposal that later brain waves like the P300 correspond to the updating, which in theory is confidence-weighted, and are enhanced by attention (Donchin, 1981; Friedman et al., 2001; Kok, 2001; Polich, 2007; Bekinschtein et al., 2009; Faugeras et al., 2012; Kolossa et al., 2013, 2015; Strauss et al., 2015). However, those later brain-waves, in particular the P300, are not systematically a signature of update, for instance in a recent EEG study (Nassar et al., 2019), the P300 was modulated by surprise, but equally and irrespective of the need to update the current estimate. In line with our results, another EEG study showed that the difference between expected and unexpected sounds (standard and deviant in a oddball task) was larger around 175-200 ms (and after 350 ms) when pupil size was larger (Hong et al., 2014). In rodents, activation of the locus coeruleus also increased the amplitude of multi-unit activity (in the sensory cortex involved in the task) specifically within 125-200 ms post-stimulus (Safaai et al., 2015). The mismatch negativity, a surprise response peaking around 170 ms, also seems better explained (in a roving paradigm) by confidence-weighted updates than by change detection, adaptation or simple prediction errors (Lieder et al., 2013). Those studies are thus consistent with the latency of the confidence-weighted surprise response we found here. Source reconstruction showed that this effect may arise in particular from the inferior frontal sulcus, which is consistent with the location of the confidence-weighted surprise responses detected with fMRI in the same probability learning task as used here (Meyniel and Dehaene, 2017).

Together, the studies quoted above and our own results support the idea that higher synchronization in alpha/beta band frequencies and lower noradrenergic activity could be mechanisms by which the brain shields current estimates against update prompted by surprising observations, thereby implementing a confidence-weighting mechanism.

## Methods

### Participants

Twenty-four participants (16 women) aged between 20 and 34 (mean: 25.4, SD: 3.7) were recruited by public advertisement. They gave their written inform consent prior to participating; the study was approved by the Ethics Committee Ile de France VII (CPP 08-021). For one subject, one of the four session was unavailable due to a technical problem.

### Task

This task and minor variants have been used previously (Meyniel et al., 2015a; Meyniel and Dehaene, 2017; Heilbron and Meyniel, 2019). The task was run using Matlab and Psychtoolbox. The experiment was divided into one training session, performed outside the MEG room, and four sessions in the MEG (lasting approximately one hour and a half with pauses). Each session presented a sequence of 380 auditory tones (one every 1.4 s), denoted A and B, corresponding to chords (350, 700 and 1400 Hz vs. 500, 1000 and 2000 Hz) that were perceived without ambiguity. Tones were presented in both ears, lasted 50 ms including 7 ms rise and fall times. The sequences were generated based on transition probabilities between successive tones; those probabilities were fixed only for a limited period of time delineated by change points. The change point probability was fixed on each trial, equal to 1/75; when a change point occurred, both transition probabilities were re-sampled within the interval 0.1 – 0.9, with the constraint that the change should be at least 4-fold for one of the probabilities. The sequence was occasionally interrupted by questions (median: every 13 trials, SD: 4.4), asking the subject to predict the identity of the next tone with a continuous slider (we reminded them about the identity of the previous tone, which is useful to report transition probabilities), and then the confidence about this prediction (slider whose ends were labeled “not at all” and “fully”).

Subjects were thoroughly instructed about this generative process. In order to acquaint subjects with the notion of randomness and transition probabilities, we used a graphical display with animated “wheels of fortune”. Each wheel corresponded to a pie chart, whose sections indicated the probability to repeat the same tone, or to change it, and each of the two charts corresponded to tones A or B, thus effectively representing transition probabilities. A ball rolled around the wheel with decaying speeding, ending at a random position (in section “repeat” or “change”) that triggered the onset of the corresponding tone. Subjects rolled the ball multiple times to generate short sequences. In order to explain the notion of change point, we introduced a key which, when pressed, changed the size of sections “repeat” and “change” randomly, in both wheels. Subjects then generated sequences with change points. We then explained to subjects that, during the task, they would only hear the sequences of tones and that they would have to figure out the underlying pie charts (i.e. transition probabilities) and the moment of change points. They performed at least one full session as training.

### Bayes-optimal learning model

The Bayes-optimal model used here is described elsewhere (Meyniel et al., 2015a, 2016; Heilbron and Meyniel, 2019) and the corresponding Matlab code is available online: https://github.com/florentmeyniel/MinimalTransitionProbsModel. We used it with the following options: the learned quantities are “transitions”, the estimation type is “HMM”, the probability grid used for numeric integration has 20*20 values and the prior about transition probabilities is flat. Below, we summarize the main aspects of this model.

The model uses Bayes rule to infer optimally the posterior distribution of transition probabilities at any given trial, denoted ***θ***_***t***_ (the bold font indicates it is a pair of transition probabilities) given a set of assumptions *M* and previous observations y_1:t_:

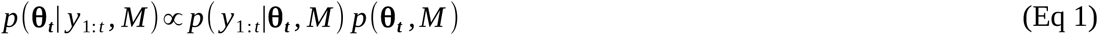

Those assumptions closely correspond to the actual generative process: the change point probability p_c_=1/75 is a given, and the model assumes that when a change point occurs, both transition probabilities are simultaneously re-sampled uniformly in the range 0 to 1 (it was not aware of additional constraints). The position of change points being unknown, the possibility that they may occur at any given trial must be taken into account. The generative process obeys the so-called Markov property: if one knows θ at time t, then the next observation y_t+1_ is generated with **θ**_**t+**1_ = **θ**_**t**_ if no change occurred and with another value drawn from the prior distribution otherwise. Therefore, if one knows **θ**_**t**_, previous observations are not needed to estimate **θ**_**t+1**_. The generative process can thus be cast as a Hidden Markov Model (HMM), which enables to iterate the computation of the joint distribution of **θ** and observations, starting from the prior, and updating this distribution by moving forward in the sequence of observations:

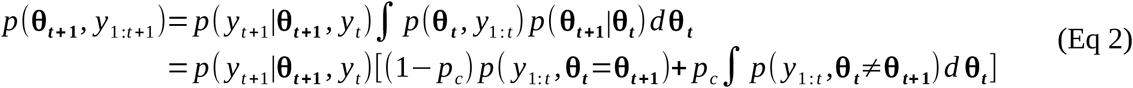

We computed this integral numerically by discretization on a grid. We obtained the posterior probability by normalizing this joint distribution.

### Formalization of prediction, surprise and confidence

The prediction, i.e. the probability of the next stimulus (question #1 asked to subjects) was computed from the posterior using Bayes rule. It is the mean of the posterior distribution of the relevant transition probability (the one that corresponds to the tone presented on the previous trial), which we note θ^rel^.

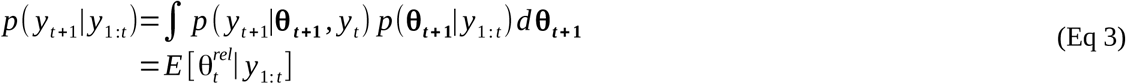

The surprise corresponding to the actual new observation was defined, following (Shannon, 1948), in bits, as the negative logarithm (to base 2) of the observation likelihood:

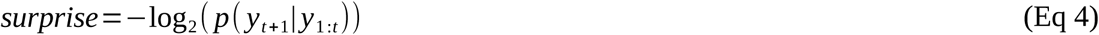

The confidence about the probability estimate (question #2) was computed as the log-precision of the posterior distribution of the relevant transition probability (Yeung and Summerfield, 2012; Meyniel et al., 2015a, 2015b; Navajas et al., 2017; Boldt et al., 2019):

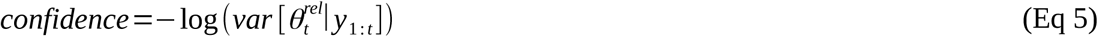

### MEG recording and pre-processing

We recorded brain activity using a whole-head MEG system (Neuromag Elekta LTD, Helsinki), sampled at 1 kHz with hardware bandpass filtering between 0.1 and 330 Hz. This systems has 102 triplets of sensors (1 magnetometer and two orthogonal planar gradiometers); we report results for the magnetometers only, but we used all sensor types for source reconstruction. We digitized the head shape (FASTRAK, Polhemus) including the nasion, pre-auricular points and various points on the scalp, and we measured head position within the MEG system at the beginning of each session with four head position indicator coils placed on the subject’s head. Electro-oculograms, electrocardiogram and pupilometry were recorded simultaneously with MEG.

Raw MEG signals were first pre-processed with the constructor software MaxFilter, in order to correct for between-session head movement (recordings realigned on the first session), removing nearby magnetic interference and correcting for noisy sensors by means of Source Space Separation (Taulu and Simola, 2006).

The data was further pre-processed with FieldTrip (Oostenveld et al., 2011). The signal was epoched from −1.4 s to 1.4 s relative to stimulus onsets, and −1.2 s to 0.5 s relative to question onsets. Epochs were visually inspected to reject those with abrupt jumps, spikes or muscular artifacts, and line filtered (50, 100, 150 Hz). Epochs were then decomposed with ICA; components were visually inspected and those resembling artifacts corresponding to eye blinks and heart beats were removed. Outlier epochs were rejected based on the signal variance and kurtosis. The mean number of trials finally included were 1335.8 ± 127 SD for stimuli and 90.2 ± 7.9 SD for questions. In order to analyze evoked responses, the signals were low-passed filtered (30 Hz) and down-sampled to 200 Hz. In order to analyze the spectral components, the signals was decomposed into peri-stimulus times (every 50 ms) and frequencies (every frequency between 7 and 40 Hz) with Morlet wavelets (using a *f/σ*_*f*_ ratio of 7, where *f* is the frequency and *σ*_*f*_ is the spectral standard deviation of the wavelet).

No baseline correction was applied since trials are not independent from one another in our transition probability task.

### Source reconstruction

We acquired anatomical T1-weighted magnetic resonance images (3T Prisma Siemens scanner) for 18 of our 24 subjects with 1 mm resolution. This anatomical image was segmented to extract the cortical surface and head shape with FreeSurfer (Dale et al., 1999; Fischl et al., 1999) and segmented tissues were imported into BrainStorm (Tadel et al., 2011) to perform source reconstruction. For subjects without a personalized anatomical image, we used BrainStorm’s default template (ICBM152). MEG and MRI data were co-registered using the digitized anatomical markers. All cortical meshes comprised ∼15 000 vertices.

We estimated at the subject-level the sources corresponding to the signal averaged within 155-230 ms (relative to stimulus onset) and across trials with high and low Bayes-optimal surprise, and high and low Bayes-optimal confidence (using median split for both). We estimated the noise covariance matrix from the signal within −0.3 to −0.1 s relative to stimulus onset, concatenated across trials. We used a normalized minimum norm estimate of the current density map, with a loose orientation constraint orthogonal to the cortical sheet (parameter 0.2), corresponding to the option dSPM in BrainStorm. The subject-level differences of current’s norm (high vs. low surprise, high vs. low confidence) where spatially normalized to the FSAverage atlas and analyzed statistically with a t-test at the group level. For display, the t-map was projected onto a high resolution mesh (∼300,000 vertices).

### Pupil size recording and pre-processing

Eye gaze and pupil size were monitored using EyeLink 1000. Due to technical errors in saving files, the data is available only for 18 of the 24 subjects. Blinks were delineated (adding a margin of 50 ms before and after) and the data within them linearly interpolated; the signal was then low-pass filtered (5 Hz) and epoched within −0.5 to 1.5 s relative to the stimulus onset. Epochs whose total blink duration exceeded 20 % were excluded. Finally, the data was z-scored per subject.

### Regression models

All multiple regression models included a constant and z-scored predictors. The significance of regression coefficients were assessed with group-level t-tests.

We often used the “residual” Bayes-optimal confidence as a predictor, which was computed by linearly regressing out the effect of prediction and predictability. More precisely, we estimated a multiple linear regression of the Bayes-optimal confidence, using as predictors the prediction itself (i.e. the Bayes-optimal estimate of the transition probability of the next stimulus, p(A)), its square and logarithm, log(p(A)) and log(p(B)). The residual Bayes-optimal confidence was defined as the residuals of this multiple regression.

The multiple regression of the evoked response shown in Fig 5 involved the post-stimulus beta-band power, testing for a relation between the evoked response and power on the same trial. Note that the sensors and timing largely overlap; a spurious correlation is thus expected due to signal expression: when recorded MEG signals from this region are stronger or less noisy (e.g. due to artifacts, sensor noise or reduced brain-sensor distance related to breathing and motion) both the measured power and evoked response should be stronger. Luckily, our prediction is that when power is stronger, the evoked response should be weaker, due to the confidence-weighting mechanism, resulting in a sign opposite to the spurious (artifactual) correlation. We predicted that if the spurious correlation is driven by fast trial-to-trial variations in signal expression (compatible e.g. with breathing), averaging successive trials into bins would disrupt it. In contrast, the expected (non spurious) correlation between power and evoked response being driven by slower trial-to-trials changes in confidence and power, it should be more robust to this binning procedure; in practice the correlation between pairs of adjacent trials was 0.61 (±0.001 s.e.m., t_23_=65.8, p=1.0 10^-27^) for confidence, and 0.15 (±0.022 s.e.m., t_23_=6.85, p=5.5 10^-7^) for power (to be more conservative, we removed linearly the effect of Bayes-optimal confidence before computing this correlation). The effect of power onto the evoked response indeed greatly increased when using non-overlapping bins of 10 consecutive trials, specifically at the moment when such a correlation is expected (around 200 ms), see **Fig S3A**. This result holds for different bin sizes (**Fig S3B**). We reproduced qualitatively this effect with simulations (**Fig S3C**) under the following assumptions: 1) beta-band power linearly reflects Bayes-optimal confidence and has auto-correlated noise from trial-to-trial, 2) the evoked response linearly reflects surprise, beta-band power (negatively) and has non-correlated noise, 3) both signals are corrupted at the measurement level by the same non-correlated noise.

### Correction for multiple comparisons

For signals with two or more dimensions (times and sensors; times, sensors and frequencies), correction for multiple comparisons was computed with FieldTrip, following permutations and cluster-based statistics (sum of t-values in the cluster) effectively controlling the family-wise error rate (Maris and Oostenveld, 2007). Sensors located less than 4 cm apart were considered as neighbors. For signals with one dimension (time), we used a custom code implementing the same cluster-based, permutation test. In the text, we systematically report the cluster-forming threshold as p-value, the cluster-level p-value and number of permutations used for the test (n).

### Cross-validated predictive accuracy

In order to estimate whether the subject’s confidence could be predicted from the pre-stimulus power across different frequencies, we used a cross-validated ridge regression. The power was averaged within −0.5 to 0 s relative to the stimulus onset and z-scored across questions. We used a ridge penalty of 0.01, but the results are quite robust to the choice of this parameter. We adopted a cross-validation approach: the data were split into 20 distinct sub-sets of inter-leaved trials and at each iteration, the ridge regression was estimated on all sub-sets but one, and its parameter estimates were used to predict subjective confidence on the left-out sub-set. We assessed the accuracy of this prediction as the Pearson correlation between predicted and actual confidence at the subject’s level.

## Acknowledgments

This work was funded by College de France and the French National Research Agency (ANR-18-CE37-0010 “CONFI-LEARN”). I thank Micha Heilbron for contributing to set up the experiment; Micha Heilbron and Maxime Maheu for recording the entire dataset and Sébastien Marti for assistance during recording.

## Supplementary figures

**Figure S1:**
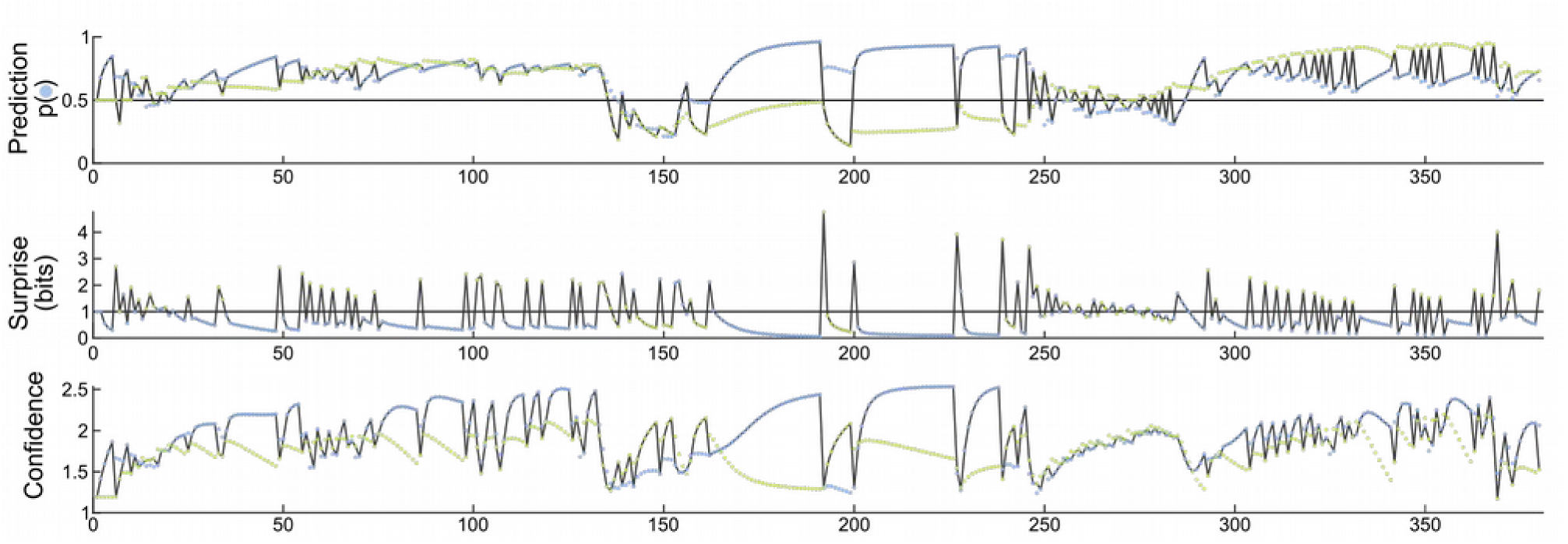
Temporal profile of confidence and surprise in the task. We consider an example sequence in which observations are color-coded (blue and green). The top graph shows the posterior inference of transition probabilities made by the Bayes optimal model in the course of sequence presentation: the green dots show the probability for the next item to be blue if the previous item was green, the blue dots show the probability for the next item to be blue if the previous item was blue. The black line shows the prediction conditioned on the identity of the item previously presented. The middle graph shows the surprise (in bits) corresponding to the actual observations, whose identity on each trial is color coded; the sequence of colored dots therefore represents the sequence of observations. The bottom graph shows the Bayes optimal confidence (i.e. posterior precision) associated with the inferred transition probabilities, using the same convention as in the top graph. Several aspects are noteworthy. First, both surprise and confidence show marked dynamics within the course of an experimental session (380 stimuli). Second, those two dynamics are distinct, for instance, surprise may be rather steady while confidence changes (e.g. from stimulus 250 to 280). Third, predictions and the associated confidence can change repeatedly from trial-to-trial when transition probabilities differ, e.g. from stimulus 300 to 340. Fourth, similar predictions can be accompanied by different confidence levels (e.g. from stimulus 100 to 140, both transition probabilities support a prediction around 0.75 and yet, the prediction is associated with higher confidence when the previous observation is blue).

**Figure S2:**
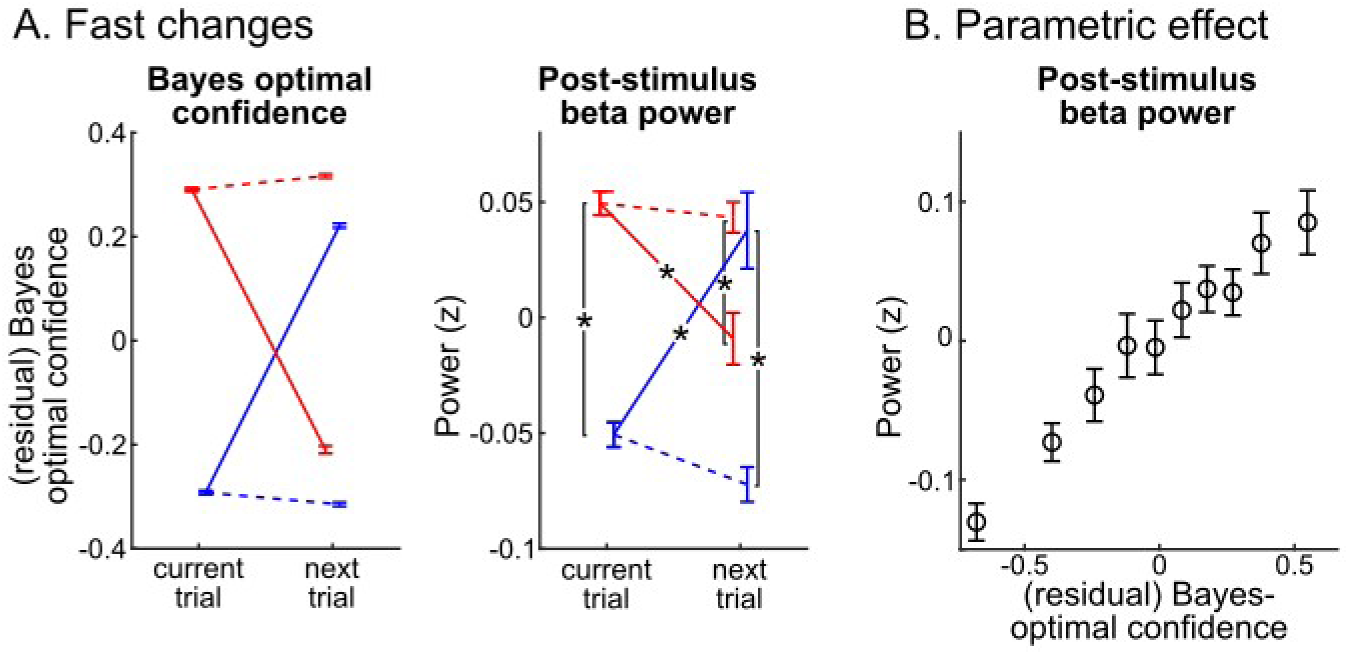
post-stimulus beta-band power shows fast changes across trials and a gradual effect of confidence. **(A)** Pairs a adjacent trials were sorted into high and low (red vs. blue) residual Bayes-optimal confidence on the current trial, and further sorted into high and low residual Bayes-optimal confidence on the next trial, therefore forming pairs that kept similar levels (dashed line) or changed drastically (plain line). The (z-scored) power in the post-stimulus beta-band cluster (Fig 3A) showed fast, trial-to-trial changes that parallel the residual Bayes-optimal confidence (*: p<0.005, paired t-test). **(B)** The correlation between post-stimulus beta-band power and (residual) Bayes-optimal confidence was not driven by specific values but indeed correspond to parametric changes. Error-bar: s.e.m.

**Figure S3:**
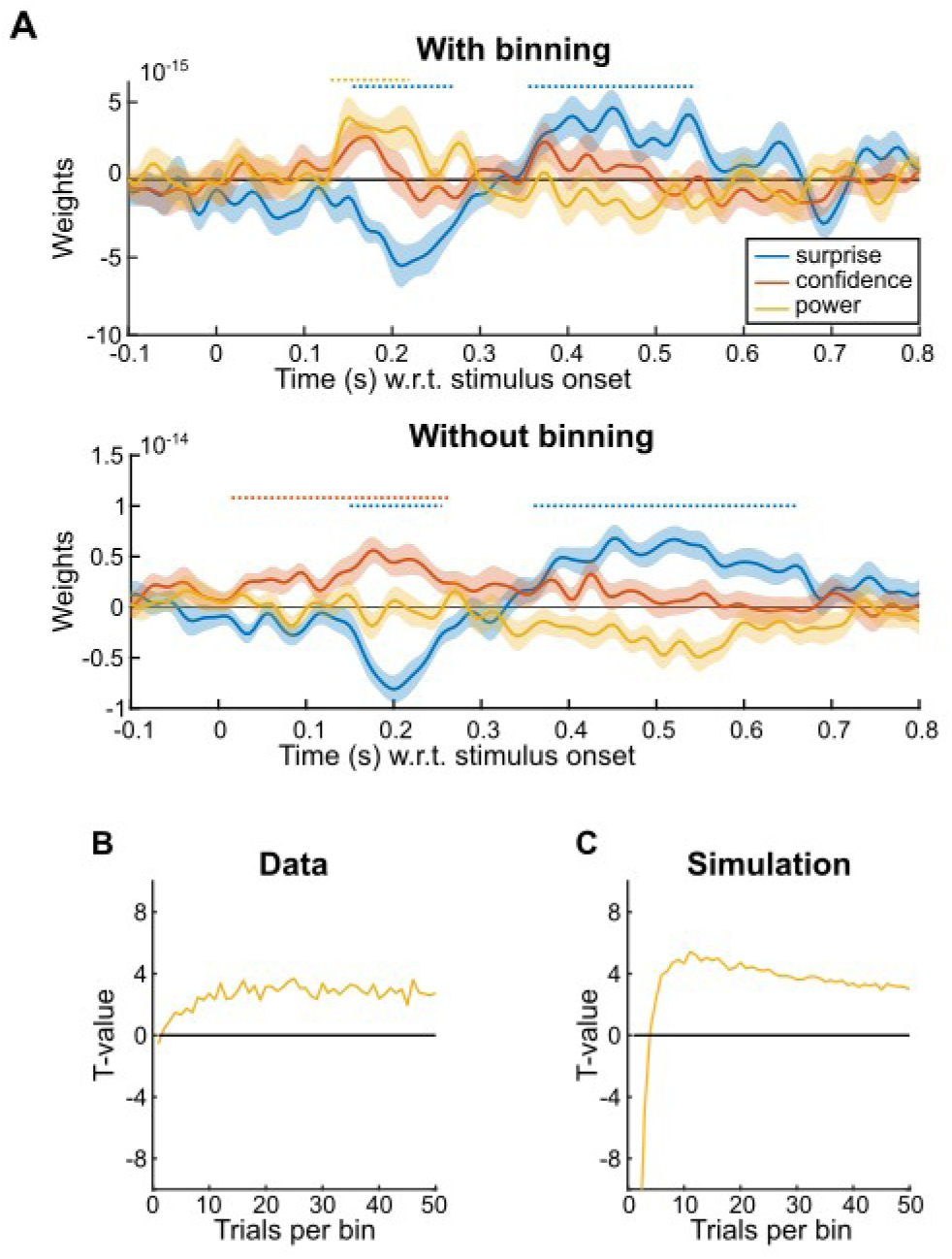
Theoretical and empirical effect of binning on the multiple regression involving power vs. evoked responses. **(A)**. The top panel is the same as Fig 5B. For this analysis, 10 consecutive trials were averaged into bins prior to estimating the regression. The bottom panel shows the results when trials are not binned (or equivalently, when there is only one trial per bin). Note the selective change around 200 ms for the effect of power. (**B)** explores this regression analysis specifically around 200 ms (averaging across significant time points, Fig 2A middle) and shows the significance (group-level t-value) of the effect of post-stimulus beta-band power onto to the ERF, depending on the number of trials per bin. (**C**) is the same analysis as in (B) but for signals simulated as follows:

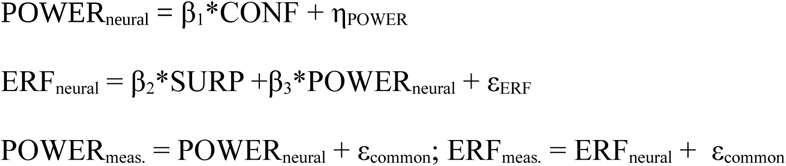

CONF and SURP are the Bayes-optimal surprise and confidence, POWER_neural_ and ERF_neural_ are the true beta-band power and evoked response, POWER_meas._ and ERF_meas._ are the signals measured by MEG; η_POWER_ is auto-correlated Gaussian noise, ε_ERF_ and ε_common_ are identically distributed Gaussian noises. For the simulation, η_POWER_ has SD=1 and auto-correlation ρ=0.5; ε_ERF_ and ε_common_ have SD=1, ρ=0; β_1_=0.25; β_2_=0.25, β_3_=0.5. Those parameters are arbitrary, not fit to the data.

